# Population structure of the invasive golden mussel (*Limnoperna fortunei*) on reservoirs from five Brazilian drainage basins

**DOI:** 10.1101/2020.05.28.122069

**Authors:** João G. R. N. Ferreira, Giordano Bruno Soares-Souza, Juliana A. Americo, Aline Dumaresq, Mauro F. Rebelo

**Affiliations:** Bio Bureau Biotechnology, Rio de Janeiro, RJ, Brazil; Institute Senai of Innovation in Biosynthetics, SENAI CETIQT, Rio de Janeiro, RJ, Brazil; Biophysics Institute Carlos Chagas Filho, Federal University of Rio de Janeiro, RJ, Brazil

## Abstract

The golden mussel (*Limnoperna fortunei*) is a freshwater bivalve that was introduced in South America almost 30 years ago, likely through ballast water of Asian ships. Since then, it has spread across the continent, causing both economic and environmental impacts. The study of the population structure of an invasive species may bring valuable insights towards understanding its pattern of dispersion, which in turn will help to create more effective management plans. Here, we have compared mussel populations from 5 different Brazilian reservoirs and tested for the presence of geographic genetic structure. In order to obtain a high number of single nucleotide variants (SNVs) at good cost-benefit, we have for the first time applied the double digest restriction-site associated DNA sequencing (ddRAD-seq) protocol for the golden mussel. The ddRAD-seq protocol allowed us to obtain 2046 SNVs, which were then used to assess population structure by applying three independent methodologies: Principal Component Analysis (PCA), Bayesian Clustering and Phylogenetic Tree. All methodologies have indicated absent geographic structure.

## 1. Introduction

Understanding the population structure of an invasive species is crucial to elucidate the demographic patterns of dispersion and to foster management plans. The golden mussel, *Limnoperna fortunei* (Dunker, 1758) is a freshwater bivalve that invaded South America at the beginning of the 1990s and since then has spread through several countries, causing both environmental and economic damage [1]. From the environmental perspective, this invasive mollusk is considered an efficient ecosystem engineer, capable of driving physicochemical changes to the water and changing the abundance of native species [2–4]. From the economic perspective, golden mussel fouling on hydroelectric plants’ equipment impacts the business by both increasing expenditure on chemicals to clean the pipes and reducing energy production due to stop for mussel-related maintenance. Only in Brazil, it is estimated that the golden mussel may cause a financial loss of up to USD 128 million a year for the electrical sector [5].

The first occurrence of *Limnoperna fortunei* in Brazil was registered almost simultaneously in two different places in 1998. One of the invasions has occurred in the Midwest region, at the State of Mato Grosso do Sul, probably from the dispersion of the population that arrived at South America in 1991 from the ballast water of Chinese ships. Another infestation event has occurred in the South region, at the Rio Grande do Sul state, probably from ballast water of ships from Argentine and/or Uruguayan ports [6]. The invasive species has then quickly spread through other states by natural or human-mediated means, the latter mainly by accidental carriage of mussels attached to ship hulls. Recently, the golden mussel has arrived in the Northeast region, on the São Francisco basin [7]. According to the collaborative database of the Center for Bioengineering of Power Plants Invasive Species, the golden mussel is present in at least 10 Brazilian states [8]. That number could increase significantly if protective policies are not taken to slow down the dispersion [9].

Population genetics has been a valuable tool for understanding the dispersal history of invasive species, typically by searching for clusters of genetically similar individuals and inferring dispersal dynamics from the identified patterns [10–12]. Interesting insights regarding the golden mussel dispersal in South America have been gained from previous studies. Zhan and colleagues have compared 22 South American populations and found support for relevant human-mediated transportation [13]. Subsequent studies have strengthened this conclusion, although the number of genetic clusters identified has differed [14,15].

In this study, we aimed at assessing the genetic structure of the golden mussel at different Brazilian reservoirs as a way to deepen our knowledge regarding the species genetic diversity and dispersal mechanisms in the country. Five different sites have been sampled: Abdon Batista, SC; Chavantes, SP; Foz do Iguaçu, PR; Três Lagoas, MS and Sobradinho, BA. Taken together, the five sampled sites represent all Brazilian regions that have golden mussel records so far: South, Southeast, Central-West, and Northeast, one of the newest infestation sites recorded so far. We have applied double digest restriction-site associated DNA sequencing (ddRAD-seq) to detect single nucleotide variants (SNVs), a strategy that allows us a better cost-benefit in the generation of data and robustness of the analyses. Different population genetic analyses (Principal Component Analysis, Bayesian clustering and Phylogenetic Tree) were done and all of them indicated absent geographic genetic structure.

## 2. Materials and Methods

### 2.1 Sample collection

The samples were collected in 5 distinct sites over the Brazilian territory (Fig. 1). The collection was done at reservoirs of hydroelectric power plants, searching for colonies of mussels attached to rocks, nets or even to power plant’s filters. Once found, the colonies were manually detached and kept (for up to 48 hours) in a plastic bucket filled with watered paper towel until they were finally stored in our lab’s aquarium.

**Fig. 1.**
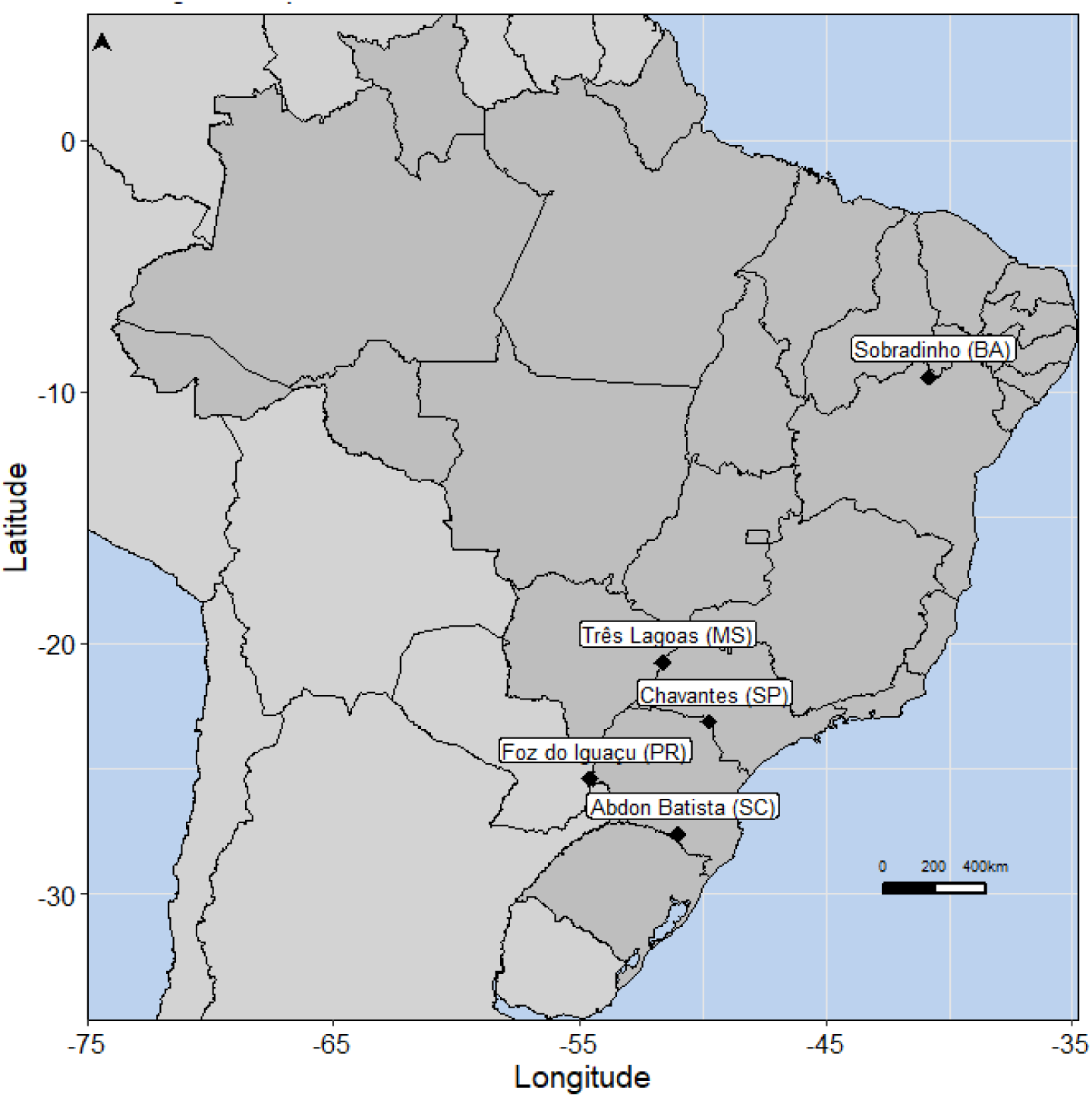
Sampling sites covered at this study, representing 5 different Brazilian states. Full name of each city is written, and the federation unit code is inside parenthesis. The geographic coordinates of each sampling sites are: Abdon Batista: 27°38’11.6”S, 51°00’03.6”W; Chavantes: 23°7’41.99”S, 49°43’59.0”W; Foz do Iguaçu: 25°26’55.9”S, 54°33’23.2”W; Sobradinho: 9°25’53.1”S, 40°49’38.7”W and Três Lagoas: 20°46’39.6”S, 51°37’44.9”W

### 2.2 DNA extraction

We have sequenced 6 mussels from each sampling site, totaling 30 mussels. Each mussel had 25 mg of its gills dissected. Finally, all of them had their DNA extracted following the protocol described by Doyle and Doyle after some adaptations [16]. Specifically, 1.5 μl of Proteinase-K (Sigma Aldrich) were added to the lysis buffer. After cell lysis, two organic extractions with 500 μl 24:1 chloroform:isoamyl alcohol solution were done, interleaved by a digestion step with 4 μl of RNAse A (Sigma Aldrich). For DNA precipitation, 0.08 volume of ammonium acetate (Vetec) 7.5 M and 0.54 volume of chilled isopropanol (Sigma Aldrich) were used.

### 2.3 ddRAD-sequencing and data pre-processing

After assessing DNA integrity, the library construction and sequencing were done by the service supplier Admera Health. The protocol applied was the same as described at the original paper that introduced the double-digest Restriction Associated DNA (ddRAD) sequencing [17]. The pair of restriction enzymes used was MluCI and NlaIII, and the pair of adapters was: adapter P1=GATCGGAAGAGCACACGTCTGAACTCCAGTCA, adapter P2=AGATCGGAAGAGCGTCGTGTAGGGAAAGAGTGT.

After library construction, DNA sequencing was done with Illumina Hiseq technology (Illumina, Inc), to generate 2×150 bp reads. The reads were demultiplexed using the software bcl2fastq (https://github.com/savytskanatalia/bcl2fastq) to separate reads by sample. Demultiplexed reads will be available upon the manuscript publication. Those reads were cleaned using the *process_radtags* module (Stacks version 2.4) with default parameters [18]. Then, *bbduk* module (BBMap version 38.69) was used to remove additional low quality bases from reads 151 bp long, converting them into 150 bp reads [19]. A quality control check of the final dataset of reads was done with FastQC version 0.11.8 [20].

### 2.4 Construction of loci catalogs

The script *stacks_denovo.pl* from the software Stacks (version 2.4) was used to build the initial catalog of loci. The script was run with parameters M=1, m=3 and n=1, after parameter optimization following Rochester and Catchen protocol [21] (Figs. S1 and S2). The *populations* function of Stacks was used for quality control, filtering out loci that had maximum observed heterozygosity higher than 90% (--max-obs-het 0.9) or minimum minor allele frequency lower than 5% (--min-maf 0.05). We have also applied a missing data filter to only allow loci that had been observed in at least 50% of samples (--min-samples-overall 0.5).

### 2.5 Population genetic analyses

The R package “hierfstat” was used to calculate population level statistics, more specifically the mean observed heterozygosity (Ho), mean expected heterozygosity (He), and the inbreeding coefficient (FIS) [22]. Pairwise FST were calculated with 10,000 bootstraps for all pairs of sampling locations using “hierfstat”‘s function *boot.ppfst*.

Principal component analysis (PCA) was applied to visualize individual relationships within and between reservoirs. The PCA was implemented using the function *dudi.pca* from the R package “adegenet” [23].

Population structure and individual admixture was inferred by applying a Bayesian clustering approach implemented on the software STRUCTURE [24]. Different numbers of assumed populations were tested (from K=1 to K=6), and for each of them 20 replicates were run. Each replica was run with 50,000 burn-in and 500,000 MCMC (Markov chain Monte Carlo) iterations. The software StrAuto (version 1.0) was used to automate the runs, as well as to calculate the most likely value of K for each catalog, according to the Evanno method [25,26]. STRUCTURE-like plots were built with scripts adapted from GitHub repository: (https://github.com/Tom-Jenkins/admixture_pie_chart_map_tutorial).

Finally, phylogenetic analysis of the samples was conducted by constructing independent trees for each locus applying the maximum-likelihood method implemented in the software RAxML (version 8.2.12) [27]. An R script (to be released upon manuscript publication) was used to screen all trees and collapse branches smaller than 0.000002 (we have identified that RAxML may create branches approximately 0.0000015 long separating identical sequences, and adding this unreal separation messed the final species tree). Then, the resulting trees were processed by the software ASTRAL (with default parameters) to estimate a single species tree [28]. FigTree (version 1.4.4) was used to create a visualization of the tree [29].

## 3. Results

### 3.1 ddRAD allows identification of 2046 SNVs and calculation of genetic population statistics

Illumina ddRADseq has yielded 214 million paired-end reads with an average of 7.2 million reads per sample. After data cleaning and quality control with process_radtags, the average number of reads per sample has decreased to 6.3 million. After application of quality control filters (as described in Materials and Methods), a final catalog was obtained containing 3316 loci, of which 2046 were variable. The catalog of variable loci was then used for subsequent population genetic analysis.

Different sampling sites have shown similar values for the mean population genetic statistics (Table 1). Observed heterozygosity ranged from 0.24 to 0.25 and it was slightly higher than gene diversity for all sites. Overall gene diversity was 0.23, which was very close to the values obtained for each site independently. Finally, inbreeding coefficient (F_IS_) was negative for all catalogs and ranged between −0.11 and −0.07.

**Table 1.** Population genetic statistics calculated for the 5 Sampling sites. Ho = mean observed heterozygosity within populations, Hs = mean gene diversity within populations; Ht = gene diversity overall; F_IS_ = inbreeding coefficient.

|  | Abdon | Três Lagoas | Sobradinho | Foz do Iguaçu | Chavantes | Average |
| --- | --- | --- | --- | --- | --- | --- |
| **Ho** | 0.24 | 0.24 | 0.25 | 0.25 | 0.25 | 0.25 |
| **Hs** | 0.22 | 0.23 | 0.23 | 0.24 | 0.23 | 0.23 |
| **$F_{IS}$** | -0.11 | -0.07 | -0.10 | -0.07 | -0.09 | -0.09 |

### 3.2 Population genetic analyses indicate absence of geographic structure

Pairwise F_ST_ values for all pairs of populations have shown little or no genetic differentiation between the sampling sites, as suggested by no 95% confidence interval greater than zero (Table 2).

**Table 2.**
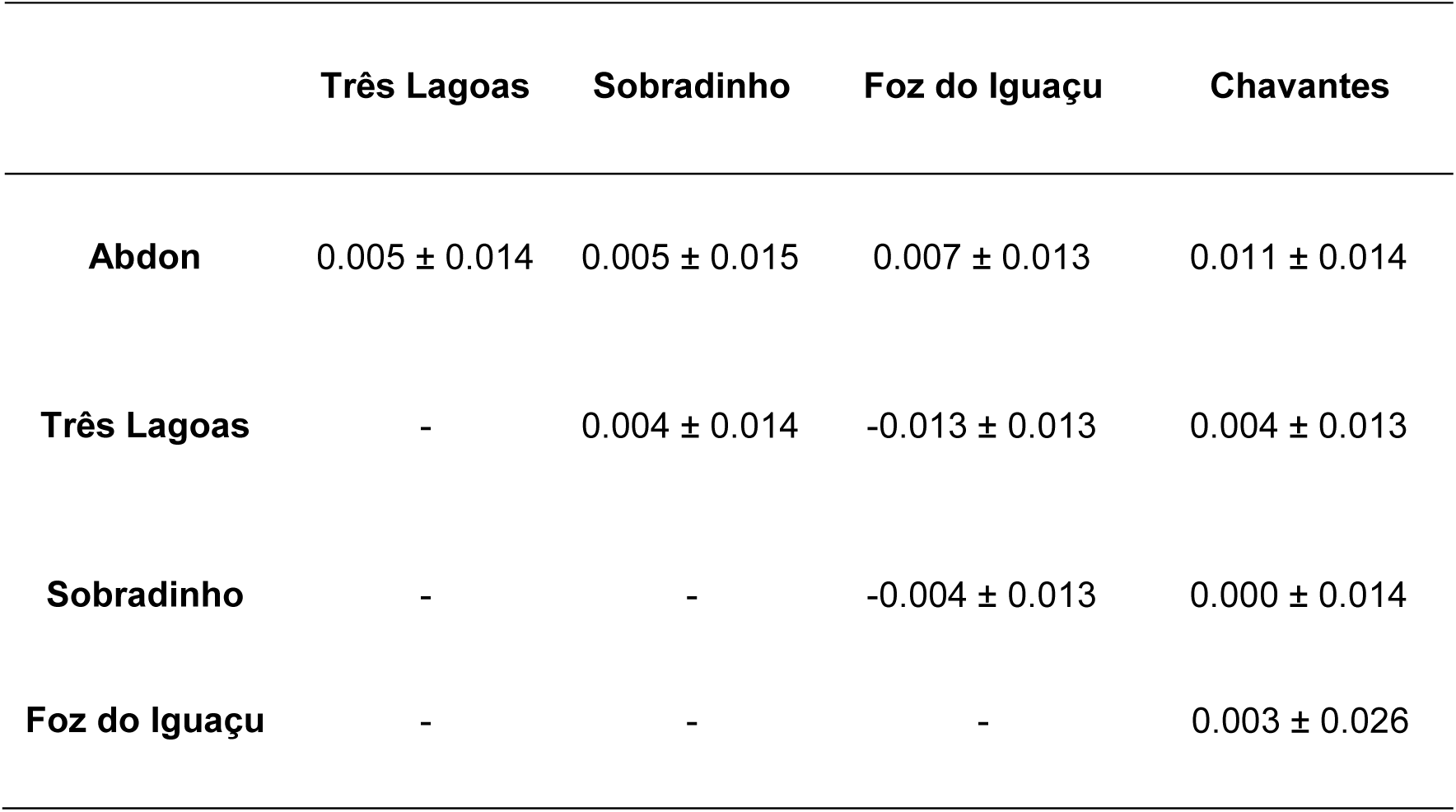
Pairwise F_ST_ values for all pairs of sampling sites. 95% confidence intervals are also displayed for each F_ST_ value.

|  | Três Lagoas | Sobradinho | Foz do Iguaçu | Chavantes |
| --- | --- | --- | --- | --- |
| **Abdon** | $0.005 \pm 0.014$ | $0.005 \pm 0.015$ | $0.007 \pm 0.013$ | $0.011 \pm 0.014$ |
| **Três Lagoas** | - | $0.004 \pm 0.014$ | $-0.013 \pm 0.013$ | $0.004 \pm 0.013$ |
| **Sobradinho** | - | - | $-0.004 \pm 0.013$ | $0.000 \pm 0.014$ |
| **Foz do Iguaçu** | - | - | - | $0.003 \pm 0.026$ |

No geographical pattern has been observed on the PCA plot (Fig. 2). Instead, the plot has shown a pattern where points (samples) from different geographic origins overlap, which could be a sign of lack of population geographic structure.

**Fig. 2.**
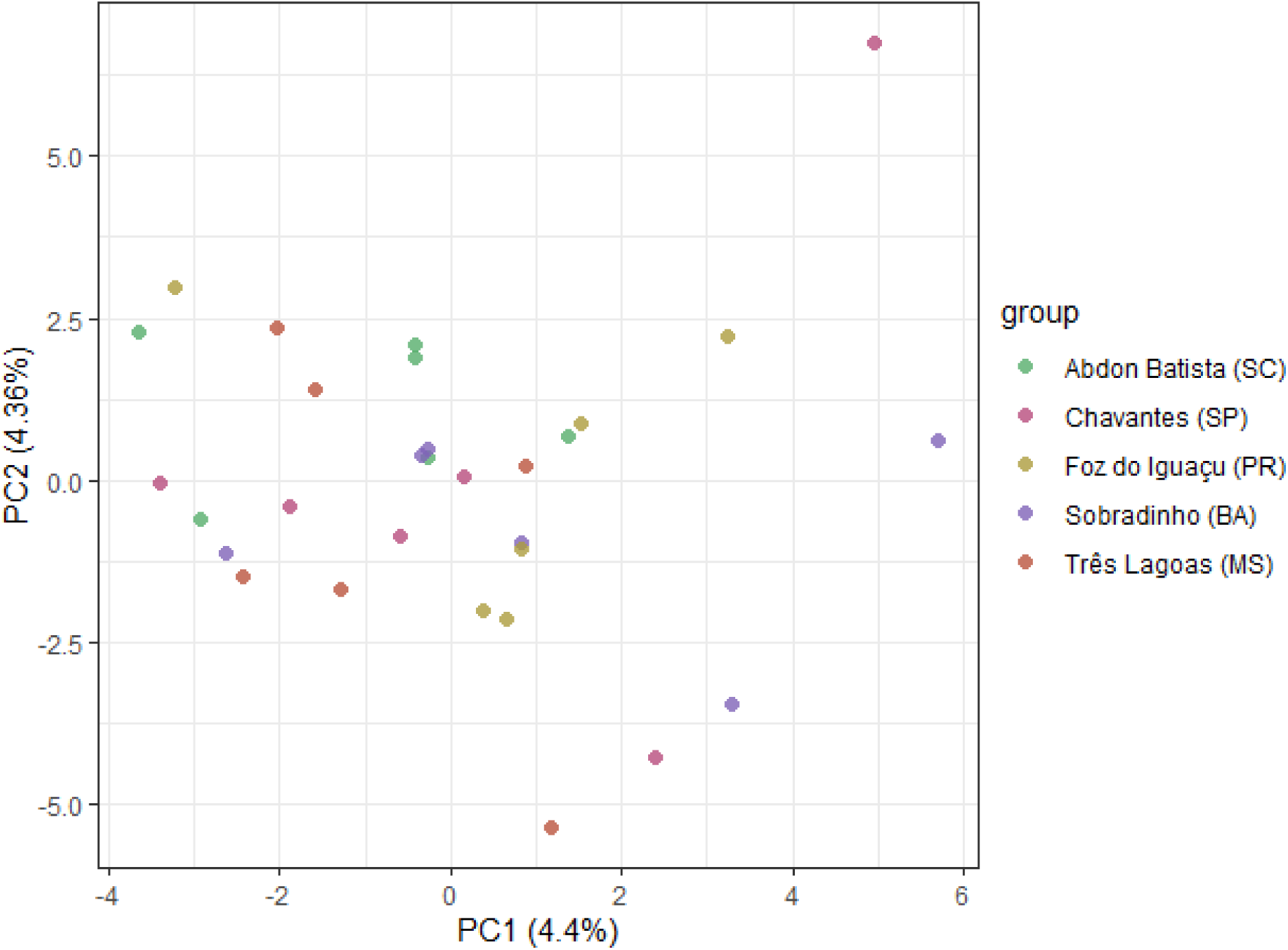
Principal component analysis (PCA) plot from the first 2 PCs. Each point represents a sample, colored according to its sampling site. Each axis (for both principal components (PC) 1 and 2) contains in parenthesis the explained variance by each PC.

The lack of geographic structure was also supported by the Bayesian clustering analysis. According to Evanno method, the most likely value for K was K=2 (by definition, the method will never return K=1 as the most likely value for K) (Fig. 3A). However, although the plot for K=2 has shown some individuals mostly belonging to one out of two possible clusters, individuals primarily from cluster 1 and 2 could be found across all sampling sites (Fig. 3B).

**Fig. 3.**
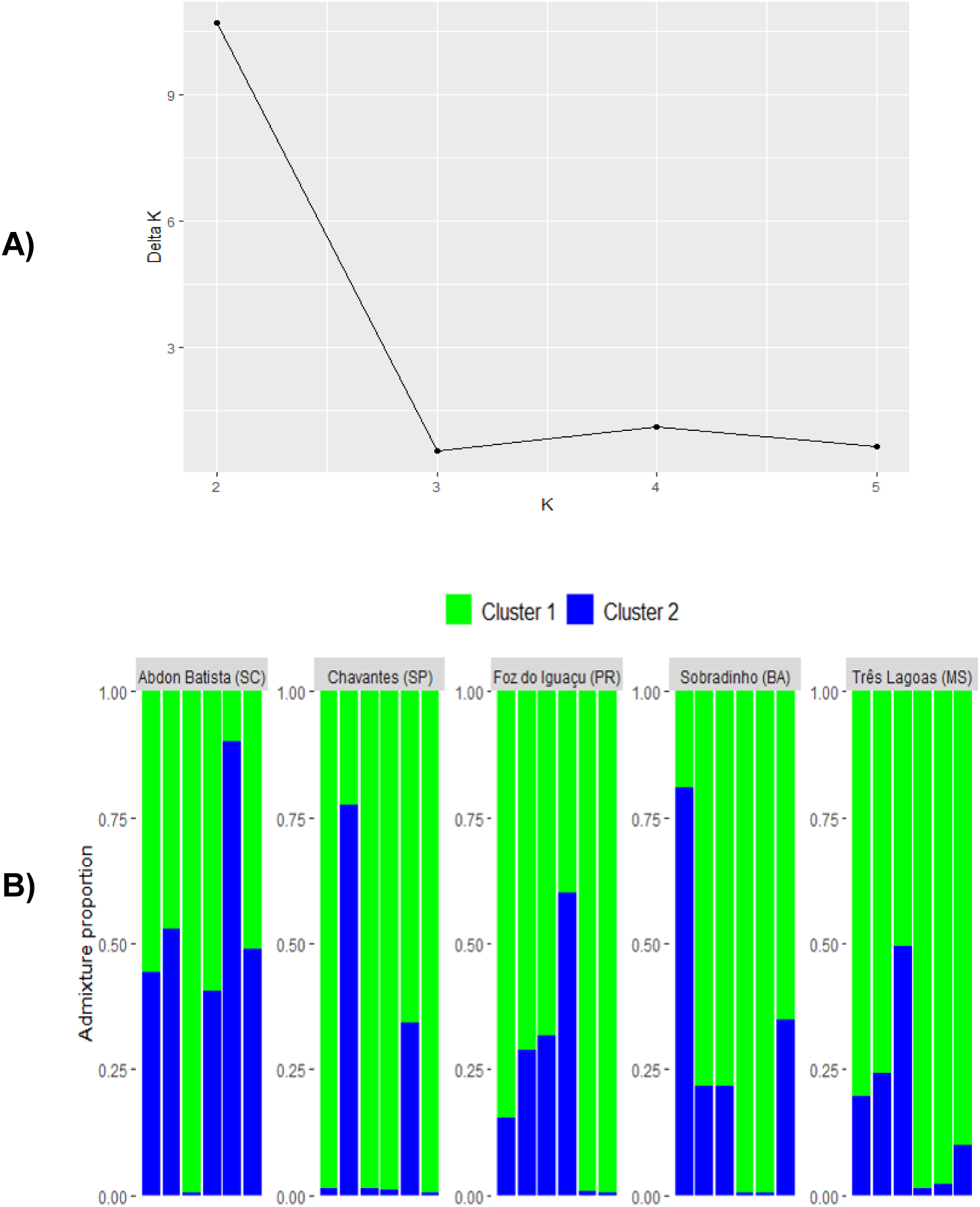
Bayesian clustering analysis. (A) DeltaK values calculated for each value of K simulated (except for the first (K=1) and last (K=6), which for definition of deltaK could not be calculated). (B) Admixture proportion estimated for each sample, considering K=2.

Finally, the phylogenetic tree has shown clades containing individuals from mixed geographic populations (Fig. 4). Besides that, most branches had low support values as calculated by ASTRAL quadripartition measurements. Taken together, those observations give stronger support for the lack of geographic structure.

**Fig. 4.**
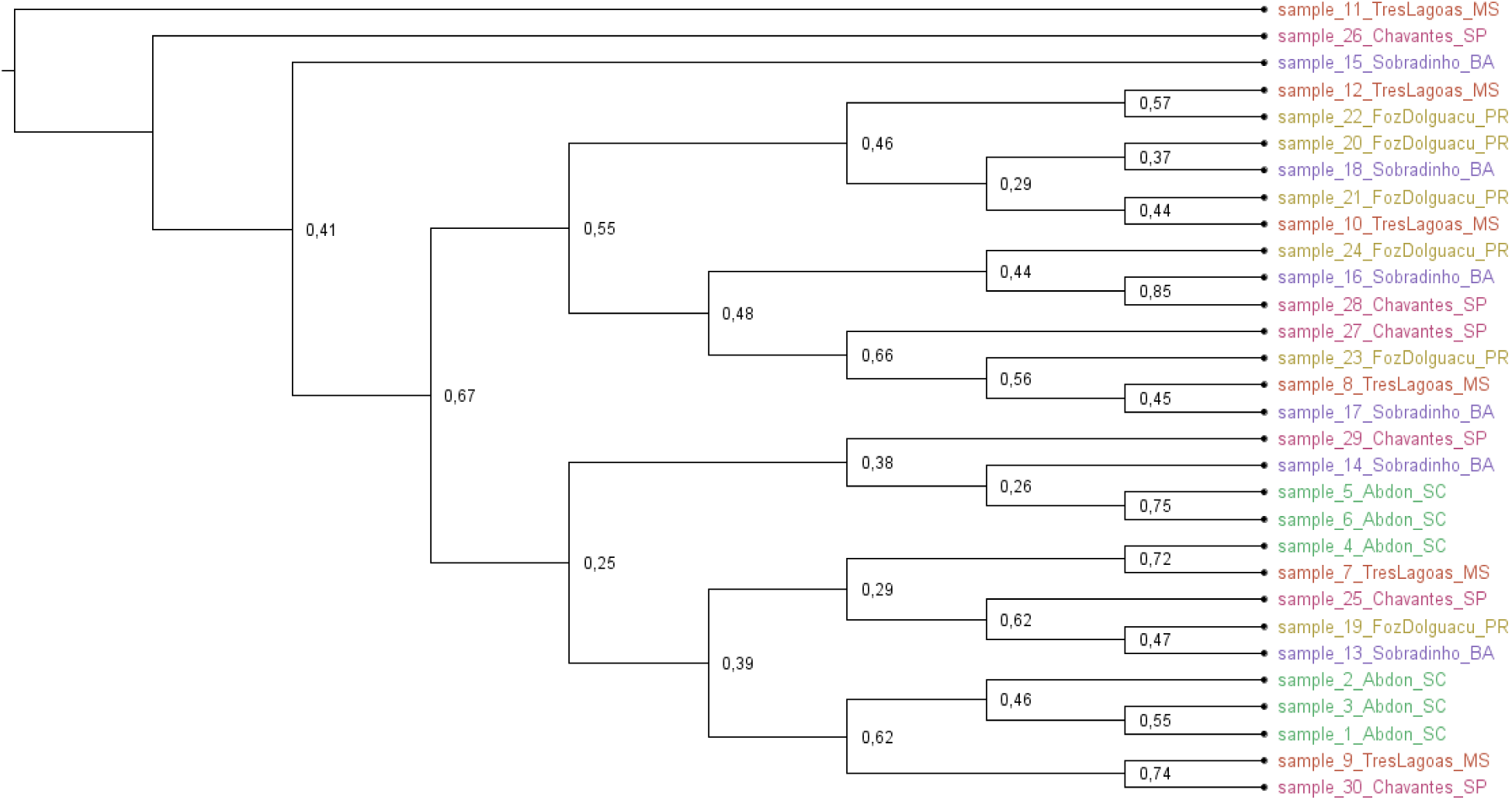
Phylogenetic trees built with ASTRAL. Posterior probabilities are displayed at each node. Each leaf represents a sample and samples are colored according to their sampling site.

## 4. Discussion

This study has not detected any evidence of geographic genetic structure between the 5 Brazilian reservoirs compared. The absence of geographic structure was independently confirmed by Principal Component Analysis, Bayesian clustering and Phylogenetic Tree. Previous studies on population genetics of the golden mussel have found similar results. For a visual summary of the Brazilian sites sampled by previous studies, refer to Fig. 5. Duarte and colleagues have analyzed 5 historically relevant South American populations using 9 allozyme loci and found no evidence that the populations were genetically differentiated [14]. The lack of geographic genetic structure was also noticed by Furlan-Murari and colleagues after sequencing 5 microsatellite loci from 3 golden mussel populations over the Paranapanema river [15]. Zhan and colleagues have sequenced 8 microsatellite loci and the mitochondrial cytochrome c oxidase subunit I (COI) gene from 22 South American populations and found 2 supposedly different genetic clusters [13]. Those clusters were not geographically separated, meaning that close populations could belong to different clusters and distant populations could belong to the same genetic cluster. This was taken as evidence supporting a jump model of dispersion, where human-mediated activities would strongly contribute to the golden mussel dispersion in the continent by jump events as opposed to gradual and slow dispersion of the species promoted by natural means. Although Zhan and colleagues have found two different genetic clusters in South America, all Brazilian populations analyzed belonged to the same cluster, i.e., no geographic structure was seen in Brazil. Nevertheless, we intend to study the effect of different filters of call rate and missing data to properly understand the impact of these parameters on the genetic structure of golden mussel.

**Fig. 5.**
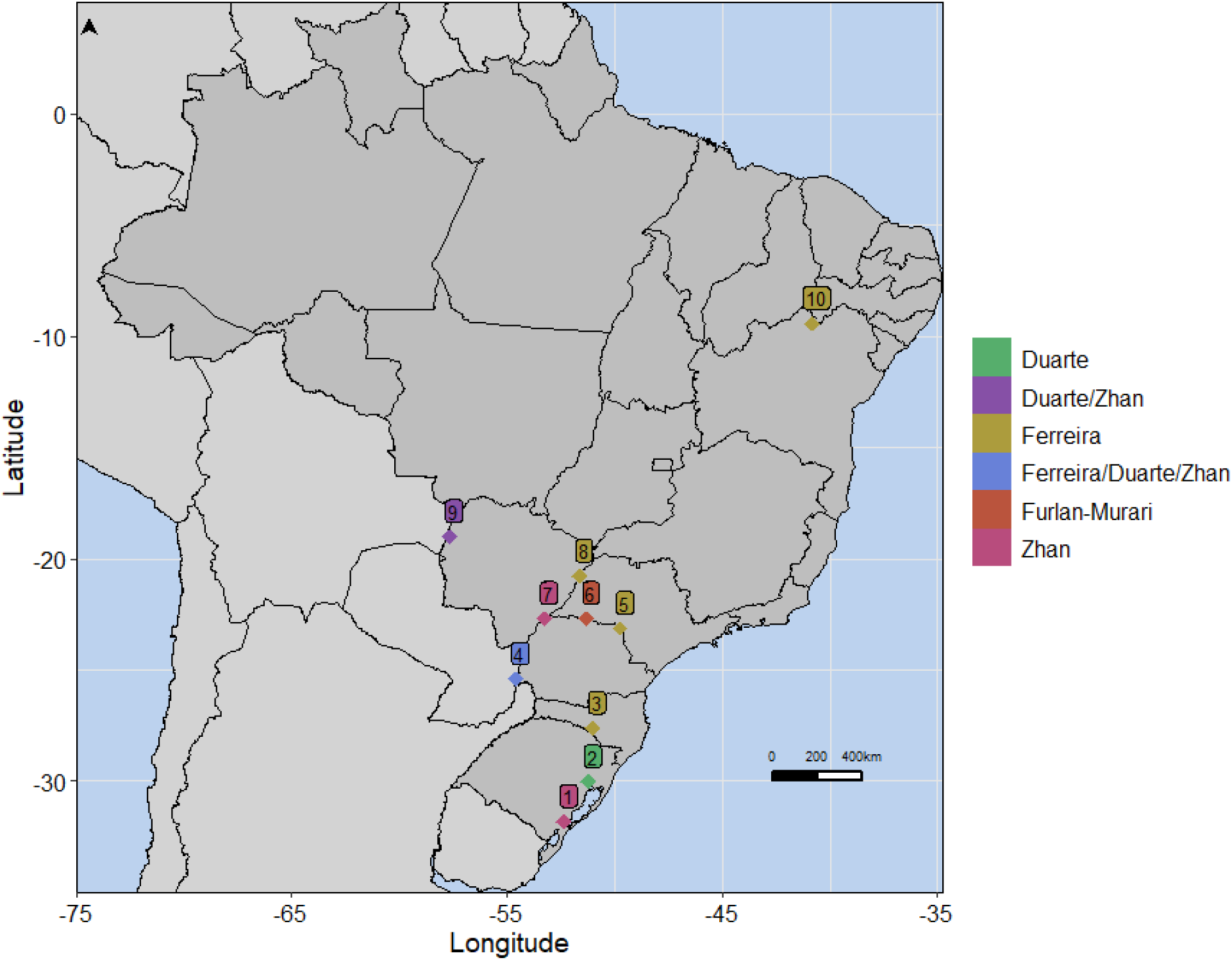
Brazilian sampling sites from studies on population genetics of the golden mussel. Different studies are colored in different ways, as well as those sites that were sampled by more than one study. Different numbers indicate different sampling sites: 1. São Gonçalo Channel, 31.81 S 52.34 W; 2. Porto Alegre, 30.03 S,51.22 W; 3. Abdon Batista, 27.61 S, 51.02 W; 4. Foz do Iguaçu, 25.41 S, 54.59 W; 5. Chavantes, 23.13 S, 49.73 W; 6. Paranapanema river, 22.69 S, 51.30 W; 7. Baía river, 22.69 S, 53.25 W; 8. Três Lagoas, 20.78 S, 51.63 W; 9. Corumbá, 19.01 S, 57.66 W; 10. Sobradinho, 9.43 S, 40.83 W.

To the best of our knowledge, this is the first time that the ddRAD protocol has been applied for the golden mussel. By applying this protocol, we were capable of retrieving 2046 variable loci, a number much higher than what was used in previous population genetic studies of the species. Although the resolution power of a single SNV marker generated by ddRAD is lower than the one from a single microsatellite, the much higher number of SNVs often obtained allows an overall greater resolution power. Previous studies have shown that several hundred SNVs are more powerful than a dozen microsatellites for standard population genetic analyses [30]. For this and other reasons like microsatellites suffering from reproducibility among laboratories, several groups have proposed to apply ddRAD (and other RAD-seq protocols) in replacement of microsatellites [18].

The lack of geographic genetic structure observed in this study supports the current hypothesis that the second infestation event in Brazil has likely occurred from ships from other South American countries as opposed to a new Asian invasion. Besides that, the lack of genetic differentiation suggests that the most recent infestation site (at Sobradinho, BA) likely originated from populations already established in South America. Given the geographic isolation of this new point, its infestation has probably arisen as a consequence of human mediated transportation, supporting the jump model of dispersion suggested by previous studies. Anthropogenic transportation inside the continent may occur by different means, such as attachment of mussels to ship hulls, discharge of ballast water, transport of alive fish in water containing mussels larvae or even commercial sand transportation [31].

Population genetics results may be of interest to other purposes besides studying the mechanisms and history of a species dispersion. Our group is currently working on a solution to fight the golden mussel invasion, and we have chosen the CRISPR/Cas9-based gene drive system as an approach for addressing the problem [5]. The basic concept is to spread across the invasive population a trait that will stop golden mussel’s reproduction. However, unforeseen mutations may compromise the effectiveness and specificity of the system. In order to develop a globally effective system, we want to choose a target site which is conserved between different individuals. Since it is not viable to sequence the target site at all living golden mussel individuals, some good criteria could be to sequence individuals belonging to different genetic clusters.

## 5. Acknowledgments

Authors are indebted to Mrs Lisboa M for her support and Wajsenzon IJR and Gondim TS for their invaluable help in experiments performance.

## 8. Supporting information captions

**S1 Fig. Number of loci shared by at least 80% of samples according to different values of parameters M and n.**

**S2 Fig. Distribution of the number of SNPS per locus, showing that the distribution pattern keeps the same over the different values of M/n parameters (M1 to M9).**

