## Supplementary figures and images for "Population structure of the invasive golden mussel (*Limnoperna fortunei*) on reservoirs from five Brazilian drainage basins"

### S1 Fig.

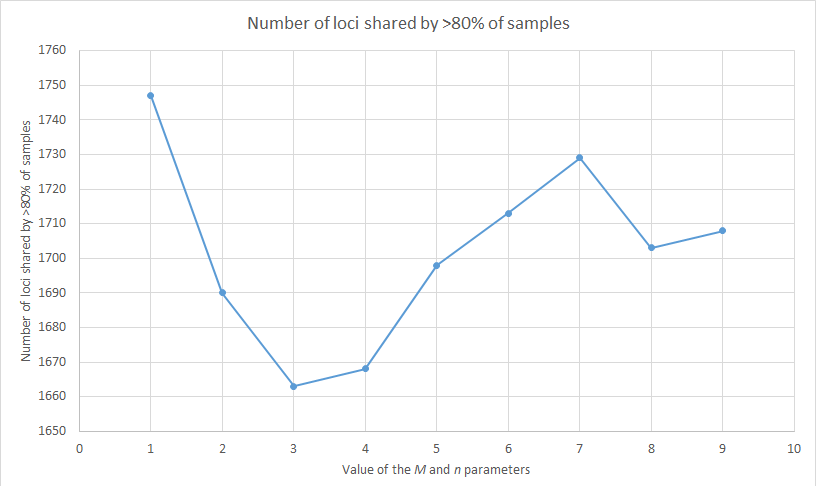

### S2 Fig.

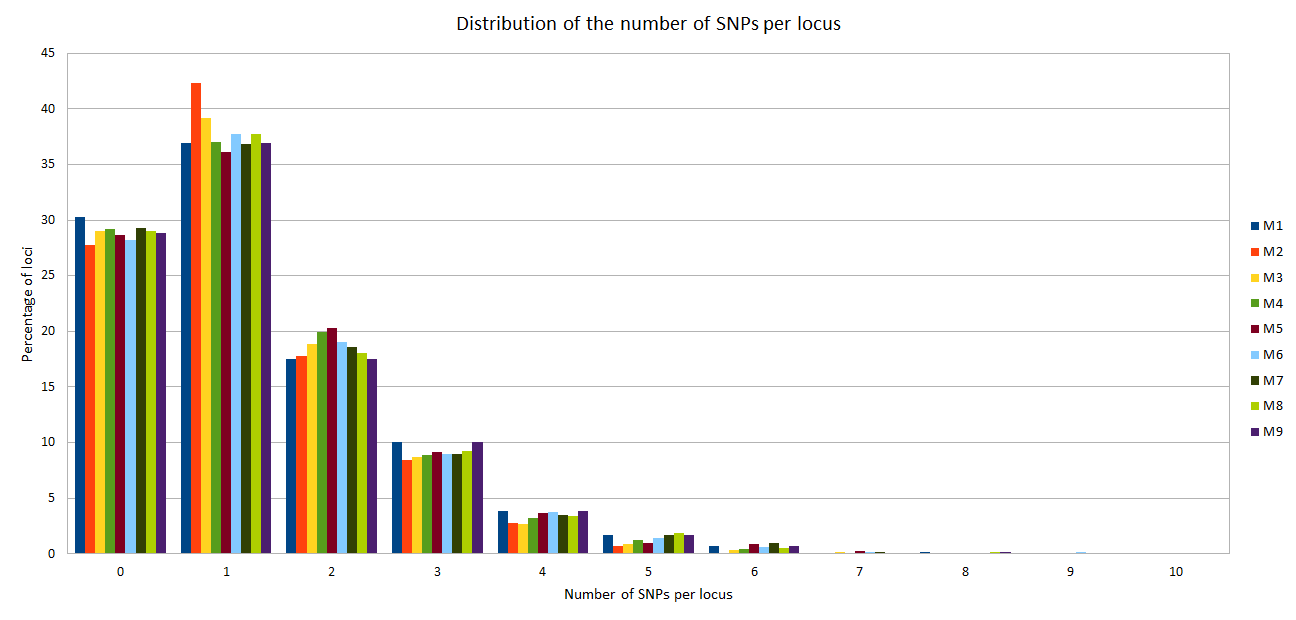
